# Modulatory Effects of Mdivi-1 on OxLDL-Induced Metabolic Alterations, Inflammatory Responses, and Foam Cell Formation in Human Monocytes

**DOI:** 10.1101/2024.12.12.628145

**Authors:** Negin Mosalmanzadeh, Rafael Moura Maurmann, Kierstin Davis, Brenda Landvoigt Schmitt, Liza Makowski, Brandt D. Pence

**Affiliations:** College of Health Sciences, University of Memphis, Memphis, TN, USA; Department of Medicine, Division of Hematology and Oncology, College of Medicine, University of Tennessee Health Science Center, Memphis, TN, USA; Center for Cancer Research, University of Tennessee Health Science Center, Memphis, TN, USA; Center for Nutraceutical and Dietary Supplement Research, University of Memphis, Memphis, TN, USA

**Keywords:** Atherosclerosis, Monocytes, Mdivi-1, Inflammation, Foam cells

## Abstract

Atherosclerosis, a major contributor to cardiovascular disease, involves lipid accumulation and inflammatory processes in arterial walls, with oxidized low-density lipoprotein (OxLDL) playing a central role. OxLDL is increased during aging and stimulates monocyte transformation into foam cells and induces metabolic reprogramming and pro-inflammatory responses, accelerating atherosclerosis progression and contributing to other age-related diseases. This study investigated the effects of Mdivi-1, a mitochondrial fission inhibitor, and S1QEL, a selective complex I-associated reactive oxygen species (ROS) inhibitor, on OxLDL-induced responses in monocytes. Healthy monocytes isolated from participants were treated with OxLDL, with or without Mdivi-1 or S1QEL, and assessed for metabolic shifts, inflammatory cytokine expression, foam cell formation, and ROS production. OxLDL treatment elevated glycolytic activity (ECAR) and expression of pro-inflammatory cytokines IL1B and CXCL8, promoting foam cell formation and mitochondrial ROS (mtROS) production. Mdivi-1 and S1QEL effectively reduced OxLDL-induced glycolytic reprogramming, inflammatory cytokine levels, and foam cell formation while limiting mtROS. These findings suggest that both Mdivi-1 and S1QEL modulate key monocyte responses to OxLDL, providing insights into potential therapeutic approaches for age-related diseases.

## Introduction

Atherosclerosis, a predominant cause of cardiovascular diseases, arises from the build-up of lipids and fibrous elements within the arteries, a process centrally involving oxidized low-density lipoprotein (OxLDL) ^1^. OxLDL is instrumental in the development of atheromatous plaques, complex arterial wall structures that are the hallmark of atherosclerosis. Infiltrating monocytes in the plaque accumulate OxLDL, evolving into foam cells ^2^. The formation of foam cells marks a critical juncture in the progression of atherosclerosis, contributing significantly to the growth and instability of arterial plaques ^3^, and foam cells increase the risk of plaque rupture which can lead to myocardial infarction or stroke ^4^.

Aging is a critical determinant in the progression of atherosclerosis, accentuating the vulnerability of arterial walls to OxLDL accumulation ^5,6^. As the body ages, the efficiency of metabolic processes and the ability to maintain homeostasis in lipid metabolism decline, facilitating the enhanced formation of OxLDL. This age-related susceptibility is compounded by increased oxidative stress and inflammatory signaling, which predispose older individuals to a higher risk of developing atherosclerotic lesions ^7^. The interplay between aging, oxidative stress, and inflammation creates a conducive environment for OxLDL-induced pathologies, underscoring the importance of investigating these processes ^5^.

OxLDL, apart from being a lipid-carrying molecule, acts as a pro-inflammatory stimulus, activating various signaling pathways within monocytes and macrophages. This activation leads to the production of inflammatory cytokines and chemokines, which further recruit immune cells to the site, exacerbating the inflammatory response. When activated by inflammation, myeloid cells undergo a significant metabolic shift towards glycolysis ^8^. Furthermore, recent studies have highlighted a critical change in cellular mitochondrial dynamics during inflammation. One of the key changes observed is an increase in mitochondrial fission, the process by which mitochondria divide ^9,10^. This increase is not just a bystander effect but is intimately linked to the metabolic reprogramming of immune cells. In this context, Mdivi-1, an inhibitor of Drp1 (Dynamin-related protein 1)-mediated mitochondrial fission, has been shown in several studies to suppress inflammatory activation in immune cells, suggesting its potential as a therapeutic agent in inflammatory and metabolic diseases ^11^.

Addressing a gap in the research, we aim to explore the modulatory effects of Mdivi-1, a pivotal regulator of mitochondrial function, on OxLDL-induced metabolic alterations, inflammatory response, and foam cell formation in human monocytes. This study seeks to broaden our understanding of how this compound can influence the progression of atherosclerosis at a cellular level, potentially offering novel insights into therapeutic strategies. We hypothesized that OxLDL would lead to glycolytic reprogramming and proinflammatory responses in monocytes, and that Mdivi-1 would reduce or prevent these responses. The implications of our research could be far-reaching, offering novel insights into therapeutic strategies that target the cellular processes underlying atherosclerosis.

## Materials and Methods

### Participants

Healthy, non-obese young adults aged 18 to 40 were recruited for the study. Participants with diagnosed conditions related to inflammation, immune dysfunction, or metabolic dysfunction, or those on medications affecting inflammation or metabolism, were excluded. All participants signed a written informed consent before participation. The study received approval from the Institutional Review Board of the University of Memphis under protocol PRO-FY2021-476. Both male and female participants were recruited without regard to race, as our previous studies showed no differential effects of race and sex on ex vivo monocyte responses to lipopolysaccharide (LPS) or SARS-CoV-2 spike protein ^12–14^, and similar results were anticipated for OxLDL in macrophages. Participants were required to visit the lab up to six times, arriving in a fasted state for blood collection for monocyte isolation. Blood was drawn through venipuncture into 1-2 10 mL EDTA vacutainer tubes, totaling 8-16 ml.

### Monocyte Isolation

Monocytes were isolated from blood samples using a negative selection method, which involved magnetic bead separation employing EasySep Direct Human Monocyte Isolation Kit (StemCell Technologies, Vancouver, BC, CAN). This technique ensured the attainment of a highly pure monocyte population with minimal contamination from other cell types. In previous studies, our methodology consistently yielded 90-95% pure monocyte populations, with no observed differences attributable to race, sex, or age ^15^.

### Metabolic Activation Assays

Isolated monocytes were seeded at 1.5 × 10^5^ cells/well in Seahorse 8-well plates (Agilent, Santa Clara, CA, USA), using Seahorse XF media enriched with 10 mM D-glucose and 2 mM L-glutamine, and total well volume was adjusted to 180 μl with supplemented media. S1QEL and Mdivi-1 (Sigma, St. Louis, MO, USA) were prepared at a stock concentration of 1 mM and stored at -20^°^C until use. Mdivi-1 was diluted to 100 μM and S1QEL was diluted to 50 μM in supplemented Seahorse media immediately before use. For Seahorse assays, 20 μl of supplemented media or inhibitor (S1QEL or Mdivi-1) were added to wells for a total volume of 200 μl. Resulting final concentrations of inhibitors were 10 μM for Mdivi-1 or 5 μM for S1QEL. The plate was centrifuged and incubated at 37°C in a non-CO_2_ incubator for one hour. OxLDL solution (2.5 mg/mL) or media was added to injection ports of the Seahorse XFp cartridge to create three groups of control (media-only), OxLDL, and OxLDL plus inhibitor. Serial measurements of oxygen consumption rate (OCR) and extracellular acidification rate (ECAR) were recorded using a Seahorse XFp analyzer (Agilent, Santa Clara, CA) for 30 min prior to and 3 hours following injection of OxLDL. Post-analysis, cell counts were conducted by photomicroscopy. Cells were subsequently lysed with TRIzol and stored at ^-^80°C for cytokine gene expression analysis.

### Inflammatory Cytokine Gene Expression

RNA was extracted from primary monocyte lysates using the TRIzol method and quantified using a Nanodrop Lite (Thermo Fisher, Waltham, MA, USA). Quantified RNA was reverse transcribed to cDNA using the High-Capacity cDNA Kit (Applied Biosystems, Waltham, MA, USA). Gene expression was analyzed by qPCR using pre-validated commercial gene expression assays and Taqman reagents (Thermo Fisher). Gene expression levels were quantified using the comparative Ct method, with ACTB as a reference gene.

### Foam Cell Assessment

Human monocytes were seeded at 2.5×10^5^ cells per well in 8-well Chamber slides, cultured in CHO-SFM II media (Gibco, Waltham, MA, USA) and treated with three conditions: media control, 1 mg/mL DiI-oxidized LDL (DiI-OxLDL), or DiI-OxLDL with 1mM Mdivi-1. Cells were incubated at 37°C with 5% CO_2_ for 48 hours before microscopic examination using a EVOS M7000 at 40x (Thermo Fisher). Foam cell formation was quantified as percentage of monocytes positive for DiI-OxLDL uptake.

### ROS Production Assessment

Isolated monocytes were plated at 2.5 × 10^5^ per well in an 8-well plate. Cells were stimulated with CHO-SFM II macrophage media (Gibco, Waltham, MA, USA) or 1 mg/mL OxLDL. After a 2.5-hour incubation, cells were washed and stained with MitoSOX™ Green (10 µM) mitochondrial superoxide indicator for 30 minutes before imaging. Cells were imaged at 40x on a fluorescence microscope (EVOS M7000, Thermo Fisher). Individual cells were quantified for fluorescence intensity in ImageJ (NIH, Bethesda, MD) and background-corrected against adjacent non-cell areas.

### Statistical Analysis

To compare two treatments, paired Student’s T-tests were employed for normally distributed data and Wilcoxon tests as a non-parametric alternative. For comparisons involving three or more treatments, repeated measures ANOVA (parametric) or Friedman tests (non-parametric) were utilized, along with Holm-Bonferroni post-hoc testing for multiple testing-adjusted mean separation. The significance level was set at α=0.05. Statistical testing was performed using R 4.4.1 (R Core Team, Vienna, AUT).

## Results

### OxLDL metabolic reprogramming and inflammation

Treatment of monocytes with OxLDL lead to an increase in acidification rate compared to the media-only control. This increase was mitigated when cells were co-treated with Mdivi-1 and OxLDL, suggesting a modulatory effect of Mdivi-1 on the glycolytic process induced by OxLDL (Figure 1A). Statistical analysis of the area under the curve (AUC) for ECAR revealed a significant increase in glycolytic activity in OxLDL-treated cells compared to control (p=0.006), and this effect was significantly reduced by Mdivi-1 co-treatment (p=0.027) (Figure 1B).

**Figure 1.**
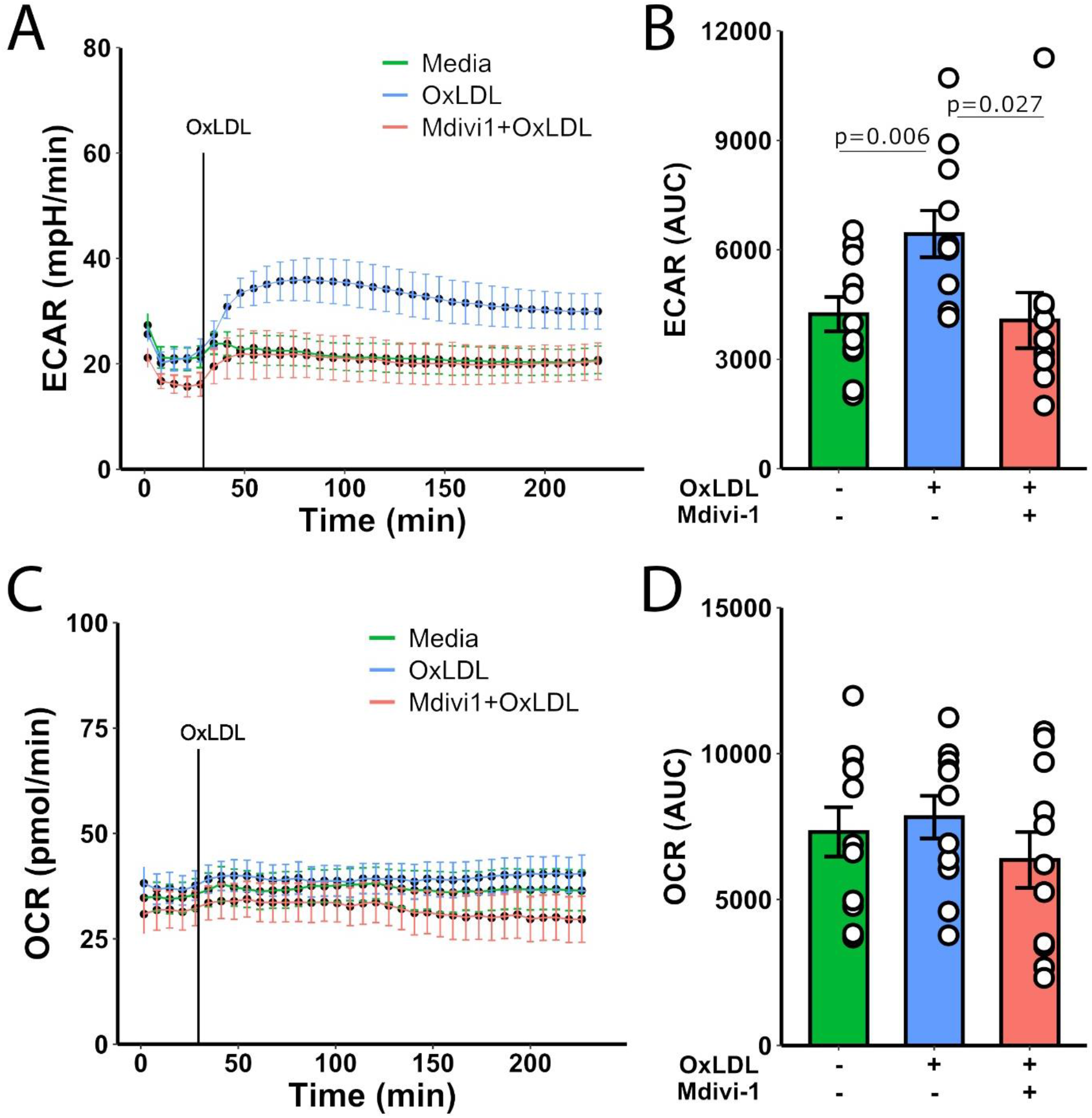
(a) Extracellular acidification rate (ECAR) measured over time in monocytes treated with media, OxLDL, or Mdivi-1+OxLDL. (b) The bar graph represents the area under the curve (AUC) for ECAR with statistical significance indicated (p=0.006 for OxLDL vs. media, p=0.027 for Mdivi-1+OxLDL vs. OxLDL). (c) Oxygen consumption rate (OCR) measured over time in monocytes treated with media, OxLDL, or Mdivi-1+OxLDL. (d) The bar graph represents the AUC for OCR.

For OCR, an indicator of mitochondrial respiration, no significant difference was observed over time across all treatment conditions (Figure 1C). The AUC analysis for OCR also indicated no significant changes, suggesting that neither OxLDL nor Mdivi-1 substantially affected mitochondrial respiration under the conditions tested (Figure 1D).

Additionally, monocyte expression of IL1B and CXCL8 was significantly elevated in the presence of OxLDL (p=0.025 and p=0.008, respectively, Figure 2A,B). Notably, the addition of Mdivi-1 to OxLDL-treated cells resulted in a significant decrease in CXCL8 expression (p<0.001), but not IL1B. IL6 and TNF expression levels did not significantly change with these treatments (Figure 2C,D).

**Figure 2.**
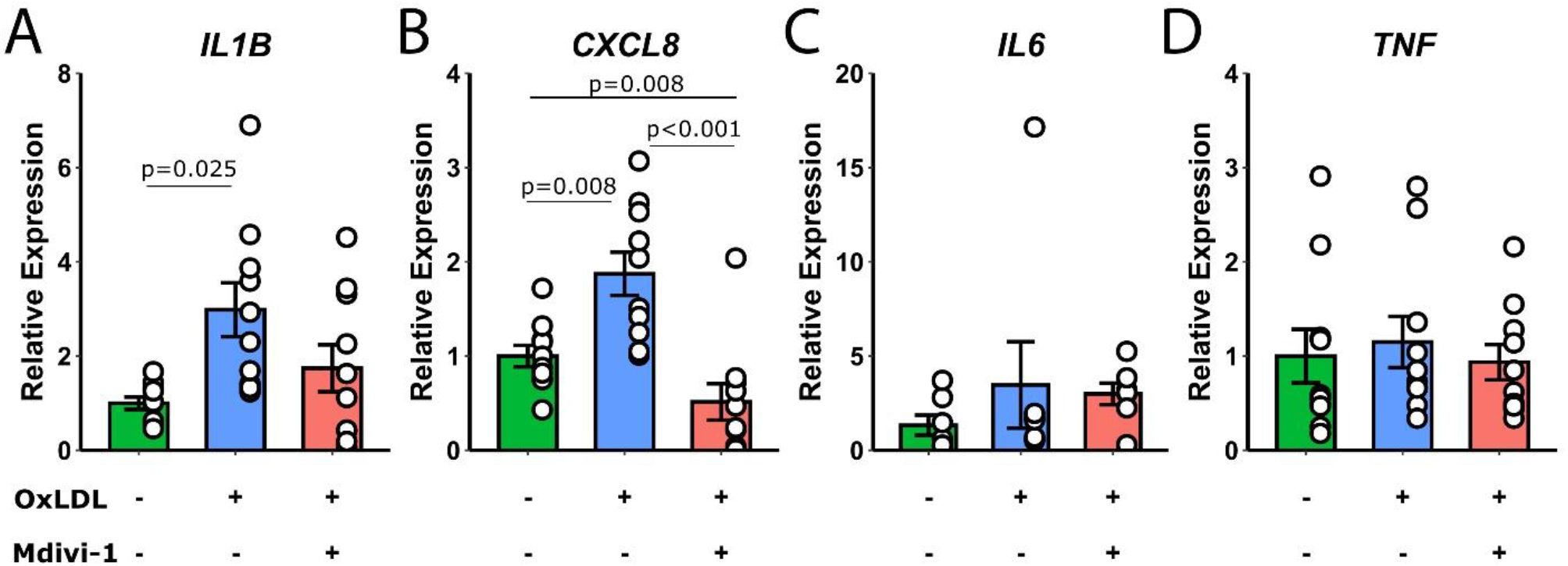
Relative expression levels of IL1B, CXCL8, IL6, and TNF in monocytes treated with media, OxLDL, or Mdivi-1+OxLDL. Statistical significance is indicated on the bar graphs.

### Foam cell formation

Foam cell formation was assessed to determine the impact of diluted DiI-OxLDL and its interaction with Mdivi-1 (Figure 3). The control group, with monocytes in standard media, exhibited no foam cell characteristics, maintaining a clear cytoplasm. Upon treatment with dil-OxLDL, a significant induction of foam cells was observed, characterized by the presence of lipid-laden vacuoles within the monocytes. The addition of Mdivi-1 to the dil-OxLDL treatment resulted in a quantifiable decrease in the number of foam cells formed (p=0.0395).

**Figure 3.**
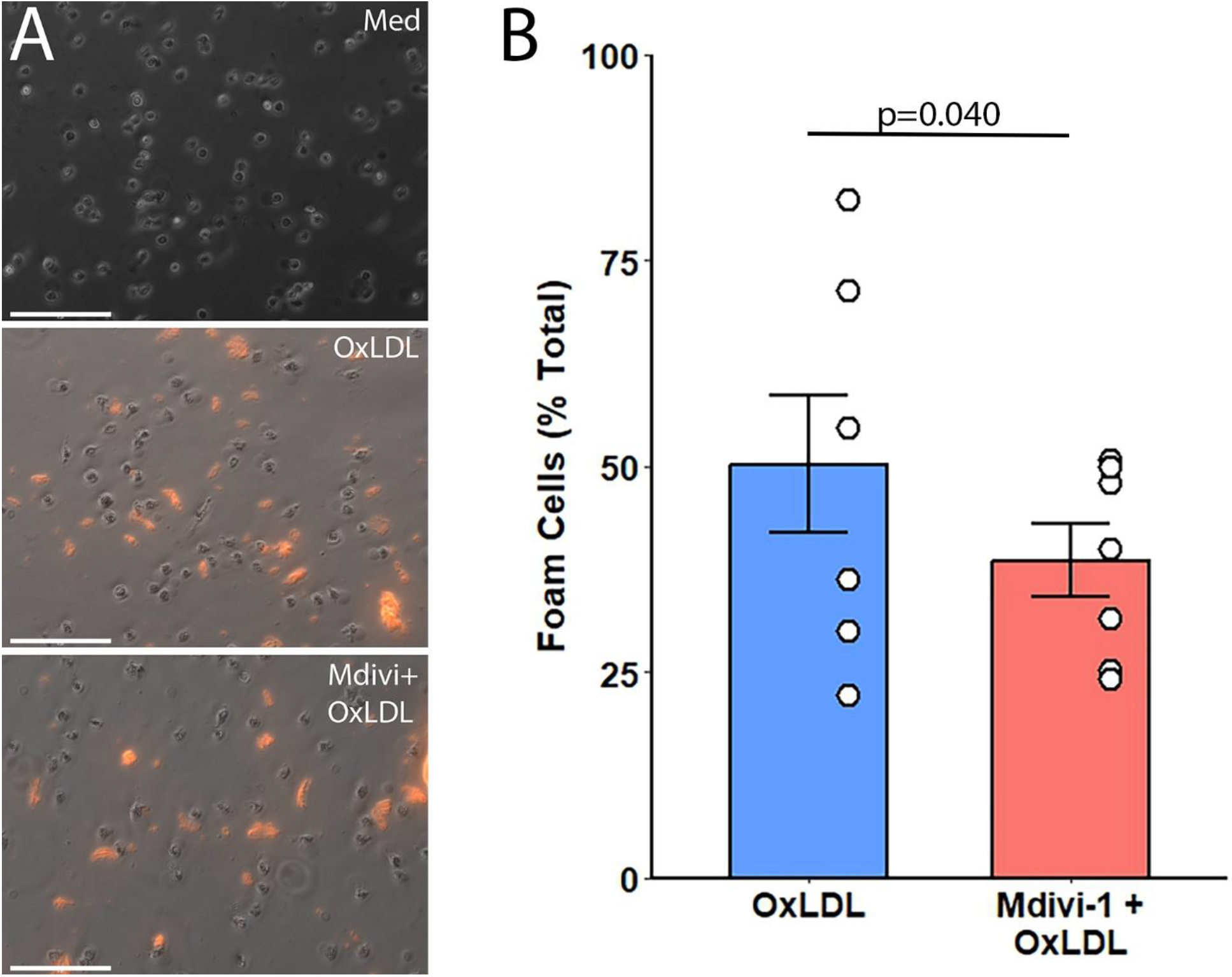
(a) Representative 40X micrographs of monocytes treated with control media, DiI-OxLDL, and Mdivi-1 + DiI-OxLDL, showing the presence of lipid-laden vacuoles indicative of foam cell formation. (b) The bar graph quantifies the percentage of foam cells in each treatment group, with statistical significance indicated (p=0.040). Scale bar = 70 µm.

The DiI-OxLDL group showed a substantial increase in the percentage of foam cells compared to the control, and the Mdivi-1 + DiI-OxLDL group demonstrated a statistically significant decrease in foam cell percentage compared to the DiI-OxLDL only group (Figure 3). This suggests that Mdivi-1 may exert a protective effect against DiI-OxLDL-induced foam cell formation in monocytes.

### Reactive oxygen species production

While Mdivi-1 is most known as an inhibitor of mitochondrial fission, it also suppresses mitochondrial reactive oxygen species (mtROS) production through mitochondrial complex I ^16^. We therefore assessed the production of mtROS in the presence or absence of OxLDL. The assessment of mtROS production revealed a notable difference between monocytes cultured in standard media and those treated with OxLDL (Figure 4). Fluorescence microscopy images showed a minimal fluorescence signal in the control group, indicating low levels of mtROS. In contrast, cells treated with OxLDL exhibited a significant increase in fluorescence intensity, indicative of elevated mtROS levels (p=0.021).

**Figure 4.**
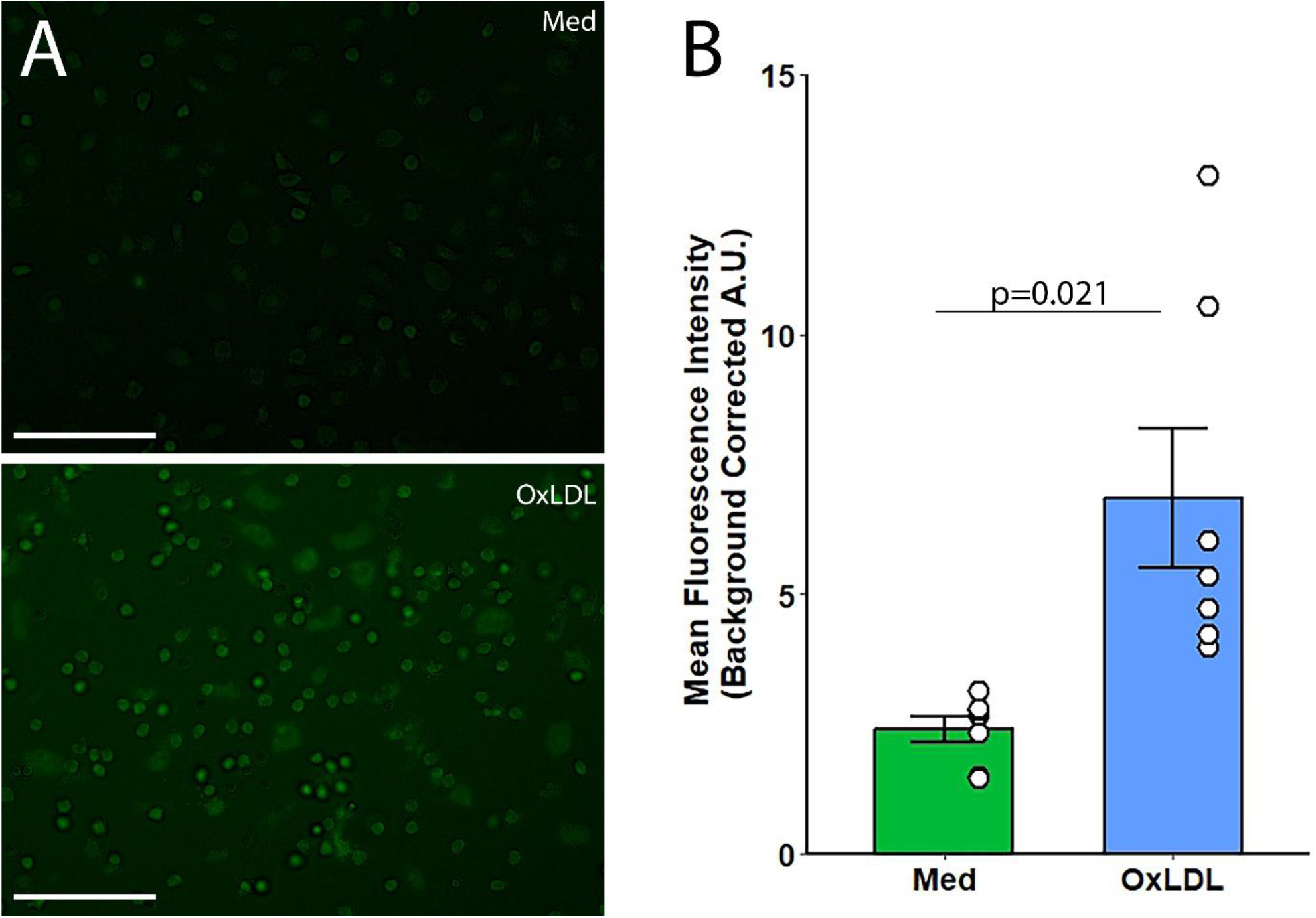
(a) Representative 40X fluorescence microscopy images of monocytes treated with OxLDL and media, showing differences in fluorescence intensity as an indicator of mtROS levels. (b) The bar graph quantifies mean fluorescence intensity (background-corrected arbitrary units, A.U.) in the different treatment groups. with statistical significance indicated (p=0.021). Scale bar = 70 µm.

A marked elevation in mean fluorescence intensity (background-corrected arbitrary units, A.U.) was present in the OxLDL-treated cells compared to the media control (Figure 4). This suggests that OxLDL treatment is associated with an increase in mitochondrial oxidative stress within monocytes

### Effects of S1QEL

Given that OxLDL increases mtROS, we examined the ability of S1QEL, a selective complex I-associated ROS inhibitor, to suppress glycolytic and inflammatory responses to OxLDL in monocytes. In general, similar trends were observed with S1QEL treatment when compared to Mdivi-1. OxLDL increased ECAR, which was significantly reduced in the presence of S1QEL (p<0.001), indicating a comparable regulatory effect of S1QEL on glycolysis (Figure 5A, B). The OCR did not significantly change with OxLDL or S1QEL treatment, suggesting that mitochondrial oxidative capacity was maintained across conditions (Figure 5C, D).

**Figure 5.**
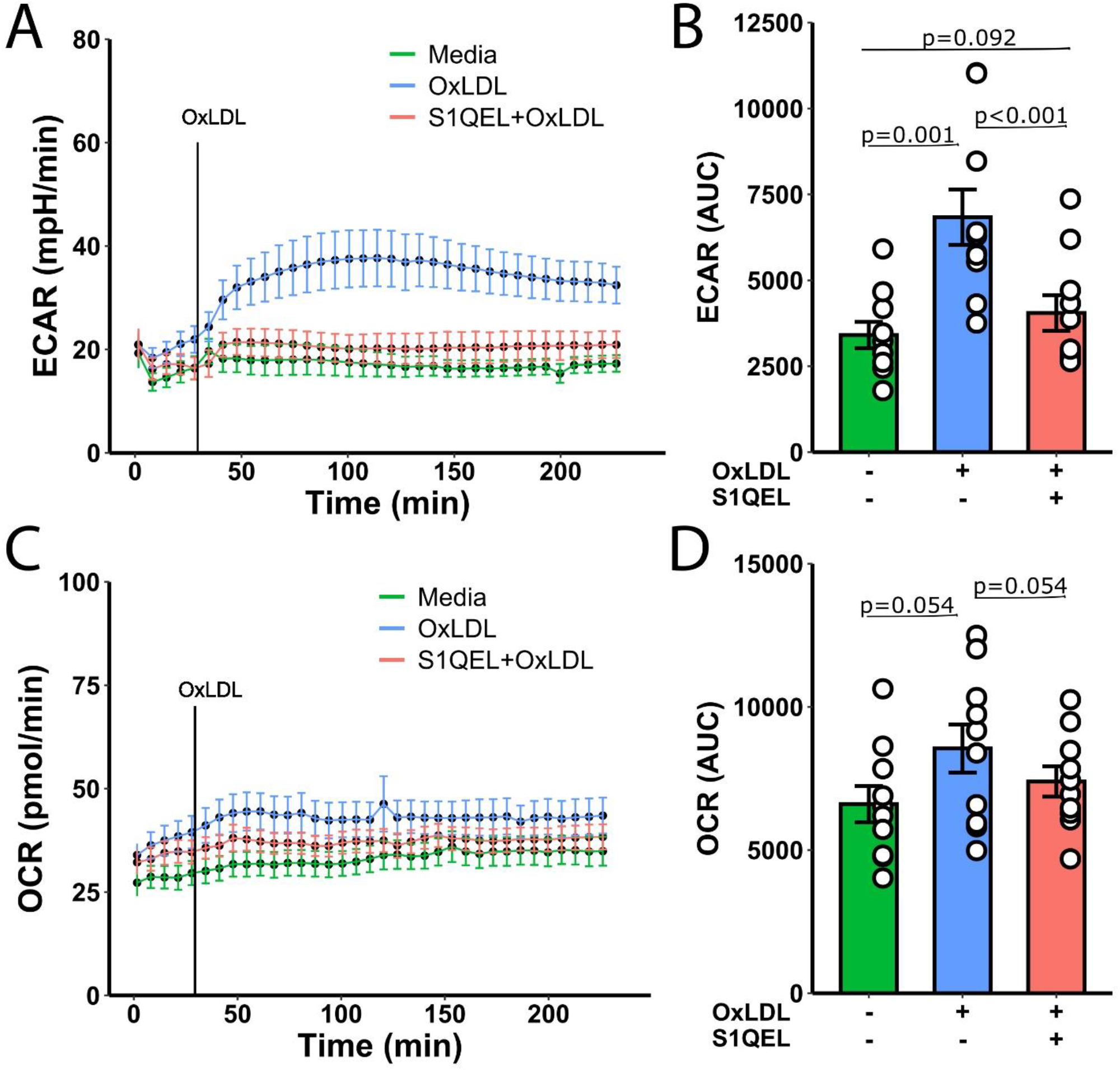
(a) Extracellular acidification rate (ECAR) measured over time in monocytes treated with media, OxLDL, or S1QEL+OxLDL. (b) The bar graph represents the AUC for ECAR with statistical significance indicated (p=0.001 for OxLDL vs. media, p<0.001 for S1QEL+OxLDL vs. OxLDL). (c) Oxygen consumption rate (OCR) measured over time in monocytes treated with media, OxLDL, or S1QEL+OxLDL. (d) The bar graph represents the AUC for OCR.

These results collectively suggest that OxLDL increases glycolytic flux in monocytes, an effect that can be modulated by both Mdivi-1 and S1QEL. However, mitochondrial respiration measured by OCR appears to be less affected by these treatments.

In Figure 6, a similar pattern was observed with the addition of the selective inhibitor of mitochondrial division, S1QEL. Treatment with OxLDL significantly increased the expression of IL1B, CXCL8, and IL6 (p=0.018, p=0.020, and p=0.003, respectively). Co-treatment with S1QEL and OxLDL significantly reduced the expression of these cytokines (IL1B: p=0.018, CXCL8: p=0.007, IL6: p=0.004). The expression of TNF remained unaltered across all conditions.

**Figure 6.**
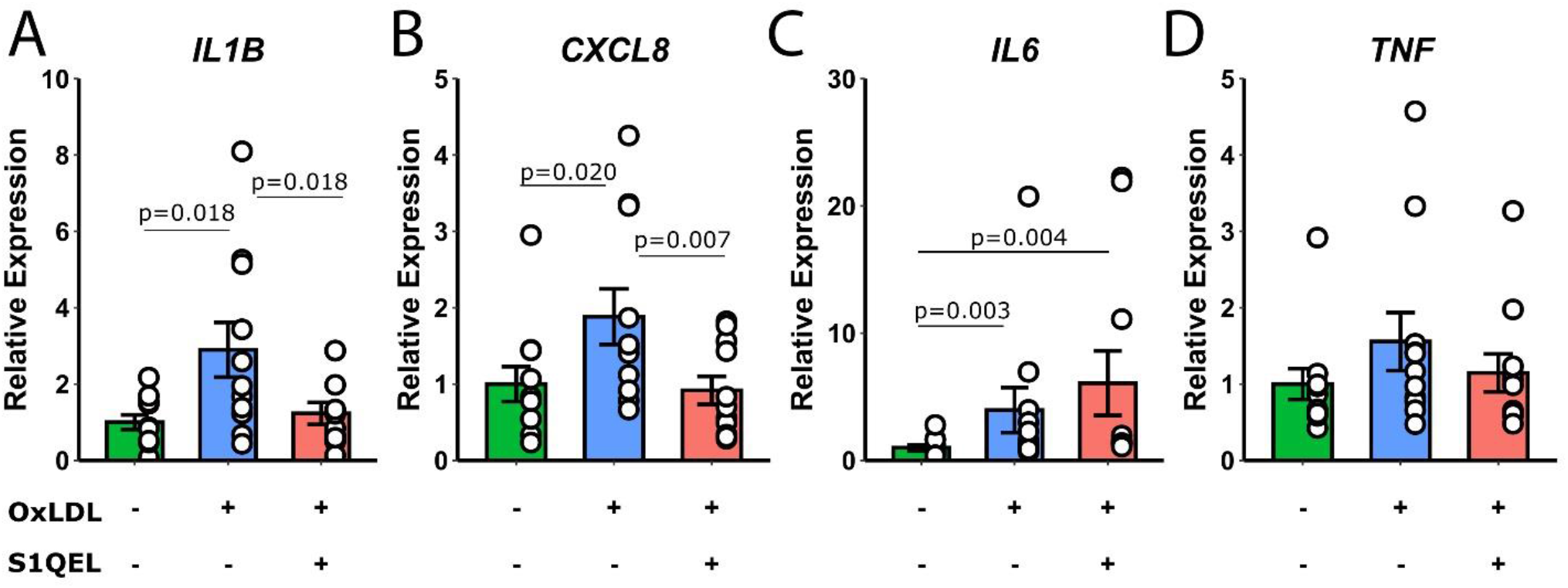
Relative expression levels of IL1B, CXCL8, IL6, and TNF in monocytes treated with media, OxLDL, or S1QEL+OxLDL. Statistical significance is indicated on the bar graphs.

These findings suggest that OxLDL can induce a pro-inflammatory state in monocytes, which is partially modulated by the mitochondrial division inhibitors Mdivi-1 and S1QEL.

## Discussion

We investigated the effects of OxLDL on monocyte glycolytic reprogramming and inflammation and assessed the potential of inhibition of mitochondrial fission using inhibitor Mdivi-1 as a strategy to ameliorate these effects. Upon exposure to OxLDL, monocytes underwent a metabolic shift toward a glycolytic phenotype, as evidenced by an increase in extracellular acidification rate (ECAR). It is likely that this is an adaptive response to meet energy needs, so that monocytes can mount oxidative stress and pro-inflammatory responses. Expression genes encoding for pro-inflammatory cytokines such as IL-1β and CXCL-8 also increased as a result of OxLDL stimulation. Our study specifically revealed that Mdivi-1 effectively mitigated the OxLDL-induced metabolic shift towards glycolysis, along with decreasing the expression of IL-1β and CXCL-8. These results highlight the potential of Mdivi-1 in counteracting the pro-inflammatory effects of OxLDL on monocytes, a key factor in the progression of atherosclerosis.

Mdivi-1 is known to inhibit Drp1, but also has been shown to suppress ROS production through reverse electron transport at mitochondrial complex I ^16^. We evaluated the ability of OxLDL to induce production of mitochondrial-derived ROS (mtROS), demonstrating an increase within the same time frame as the inflammatory and glycolytic responses noted in our initial experiments. To determine if Mdivi-1 might be acting through inhibiting mtROS production, we evaluated the complex I ROS-specific inhibitor S1QEL ^17^. S1QEL pretreatment had very similar effects to Mdivi-1, including suppressing glycolytic reprogramming and altering expression of *IL1B* and *CXCL8*, the same genes inhibited by Mdivi-1. This suggests that Mdivi-1 is likely mediating anti-inflammatory effects through suppression of complex I-specific mtROS production.

In addition to these metabolic and inflammatory changes, OxLDL leads to excessive lipid internalization, driving the transformation of these cells into foam cells. These foam cells, characterized by large lipid droplets, are a hallmark of atherosclerotic plaques. As these foam cells accumulate within the arterial wall, they disrupt the normal structure and function of the vasculature, contributing to the thickening and stiffening of arterial walls. Our findings support and extend previous research on Mdivi-1 effects on cellular responses related to atherosclerosis in several other cell types. Fang et al. focused on the protective effects of inhibiting mitochondrial fission on OxLDL-induced vascular smooth muscle cell (VSMC) foaming. Mdivi-1 antagonized the damaging effects of OxLDL on mitochondrial morphology and reduced foam cell formation in VSMCs, highlighting its potential in the treatment of atherosclerosis ^18^. This evidence aligns with our findings in monocytes, suggesting a broader applicability of Mdivi-1 in combating OxLDL-induced cellular damage. Moreover, research by Ze-da-Zhong Su et al. (2023) demonstrated that Mdivi-1 reduced plaque areas, M1-like macrophage polarization, NLRP3 inflammasome activation, and Drp1 phosphorylation. These results not only reinforce the anti-atherosclerotic potential of Mdivi-1 but also its ability to influence key inflammatory pathways and macrophage behavior, crucial in atherosclerosis pathogenesis ^19^.

Furthermore, the anti-inflammatory properties of Mdivi-1 were highlighted in a study by Li et al., which examined its effects on atopic dermatitis (AD) ^20^. In this study, human keratinocytes pre-treated with Mdivi-1 and stimulated with an inflammatory cocktail exhibited reduced symptoms of AD. The study concluded that Mdivi-1 alleviates AD symptoms by inhibiting NLRP3 inflammasome activation and blocking the NFκB pathway. This finding is particularly relevant as it underscores the broader anti-inflammatory and immunomodulatory capacities of Mdivi-1, beyond its role in atherosclerosis. In light of these studies, it is evident that Mdivi-1 not only mitigates the effects of OxLDL in monocytes, as demonstrated in our findings, but also exerts a broader spectrum of protective effects in different cell types and disease contexts. This versatility enhances the therapeutic potential of Mdivi-1, making it a promising candidate for further research and clinical application in treating atherosclerosis and other inflammatory diseases.

Alongside the significant effects of Mdivi-1, our study also investigated the impact of S1QEL on monocyte function in the context of OxLDL exposure. We observed that S1QEL, much like Mdivi-1, played a pivotal role in modulating the metabolic and inflammatory responses induced by OxLDL. S1QEL effectively suppressed the glycolytic shift and inflammatory cytokine expression in monocytes, suggesting its potential as a therapeutic agent in atherosclerosis.

There is limited literature on the effects of S1QEL on inflammatory responses. Sanchez-Perez in 2020 focused on its application in myocardial ischemia-reperfusion (IR) injury, demonstrating that S1QEL significantly reduced infarct size in mice lacking mitochondrial uncoupling protein 3 (UCP3), indicating its crucial role in contexts where exacerbated superoxide production is a concern ^21^. Notably, the absence of UCP3 led to significant mitochondrial structural changes, particularly in older mice, underscoring the potential of S1QEL as a therapeutic strategy in mitochondrial-related cardiac diseases. This suggests that S1QEL, through its unique action on mitochondrial complex I, can be a potent agent in addressing not only the metabolic and inflammatory changes in monocytes induced by OxLDL (as in our study), but also in broader contexts of mitochondrial dysfunction and oxidative stress. The findings from our study, in conjunction with the existing literature, reinforce the potential of S1QEL as a promising therapeutic strategy, not only in the context of atherosclerosis but also in other conditions characterized by mitochondrial impairment and oxidative stress.

The present study bridges a significant gap in the current literature, particularly in the context of Mdivi-1 and S1QEL’s effects on monocytes within atherosclerosis. Notably, while the role of Mdivi-1 has been somewhat explored in vascular smooth muscle cells and macrophages, its specific impact on monocytes, especially under OxLDL influence, has not been previously examined. Additionally, the literature on S1QEL is remarkably limited. We provide key insights into how both Mdivi-1 and S1QEL modulate metabolic and inflammatory responses in monocytes, an essential factor in the progression of atherosclerosis and crucial given the pivotal role of monocytes in the development of atherosclerotic lesions.

While our findings are promising, they have several limitations. Chiefly, *in vitro* results may not completely represent the outcomes in a more complex *in vivo* setting. Cellular interactions and systemic physiological responses in a living organism can significantly influence the effectiveness of therapeutic agents like Mdivi-1 and S1QEL. Further studies, particularly those involving animal models or clinical trials, are necessary to validate and extend our findings. Moreover, while our study focuses on atherosclerosis, the mechanisms uncovered may have implications for other conditions characterized by oxidative stress and inflammation. Exploring the potential of Mdivi-1 and S1QEL in these broader contexts could open new avenues for therapeutic interventions in various diseases. Additionally, our research did not address the long-term effects of Mdivi-1 and S1QEL treatment, which are critical for assessing their sustained impact and potential side effects. This gap highlights the need for extended studies and clinical trials to evaluate the long-term efficacy and safety of these compounds, thereby establishing their clinical relevance and therapeutic potential in the treatment of atherosclerosis and related cardiovascular diseases.

## Conclusion

In conclusion, our study demonstrates the therapeutic potential of Mdivi-1 and S1QEL in countering pro-inflammatory monocyte responses to OxLDL, a key step in the development of atherosclerosis. We demonstrated that Mdivi-1 effectively reduces the metabolic shift towards glycolysis induced by OxLDL, decreases foam cell formation, and lowers gene expression of inflammatory cytokines like IL-1β and CXCL8. Treatment with the complex I-specific mtROS inhibitor S1QEL similarly modulates the same metabolic and inflammatory responses to OxLDL in monocytes. These findings not only advance our understanding of the disease’s pathophysiology but also pave the way for future *in vivo* studies and the exploration of potential broader therapeutic applications (beyond atherosclerosis) of these compounds.

## Availability of data and materials

The datasets and analytical scripts used in preparing this manuscript are available in the FigShare repository (https://doi.org/10.6084/m9.figshare.c.7498686). *AUTHOR NOTE: this link will not be active until the FigShare repository is published. Once published, the FigShare repository can no longer be edited or added to, so this will be done upon acceptance of the manuscript. We are happy to provide access to datasets on reviewer request*.

## Acknowledgements

The authors thank the study participants for their contributions.

## Ethics declarations

### Ethics approval and consent to participate

The study received approval from the Institutional Review Board of the University of Memphis under protocol PRO-FY2021-476. All participants provided written informed consent prior to enrolling in the study.

### Consent for publication

Not applicable

